# Defective phagocytosis leads to neurodegeneration through systemic increased innate immune signaling

**DOI:** 10.1101/2023.01.08.523170

**Authors:** Johnny E. Elguero, Guangmei Liu, Katherine Tiemeyer, Heena Gandevia, Lauren Duro, Kimberly McCall

## Abstract

In nervous system development, disease and injury, neurons undergo programmed cell death, leaving behind cell corpses that are removed by phagocytic glia. Altered glial phagocytosis has been implicated in several neurological diseases including Alzheimer’s disease, Parkinson’s disease, and traumatic brain injury. To untangle the links between glial phagocytosis and neurodegeneration, we investigated *Drosophila* mutants lacking the phagocytic receptor Draper. Loss of Draper leads to persistent neuronal cell corpses and age-dependent neurodegeneration. Here we investigate whether the phagocytic defects observed in *draper* mutants lead to chronic increased immune activation that promotes neurodegeneration. A major immune response in *Drosophila* is the activation of two NFκB signaling pathways that produce antimicrobial peptides, primarily in the fat body. We found that the antimicrobial peptide Attacin-A is highly upregulated in the fat body of aged *draper* mutants and that inhibition of the Immune deficiency (Imd) pathway in the glia and fat body of *draper* mutants led to reduced neurodegeneration, indicating that immune activation promotes neurodegeneration in *draper* mutants. Taken together, these findings indicate that phagocytic defects lead to neurodegeneration via increased immune signaling, both systemically and locally in the brain.

## Introduction

The link between neurodegeneration and innate immunity has been the subject of much research in recent years ^1–4^. Innate immunity consists of two main branches: the cellular branch, or phagocytosis, and the humoral branch, which is comprised of signaling pathways culminating in the release of cytokines, antimicrobial peptides (AMPs), and other immune molecules ^5,6^. One aspect of innate immunity, inflammation, can be defined as the activation of immune cells coupled with the release of inflammatory cytokines. This response is important for the protection of the organism from pathogens as well as for tissue repair after injury. Although inflammation is protective in the context of infection and injury, chronic inflammation is known to be detrimental ^7,8^. The presence of inflammation in the central nervous system has been termed “neuroinflammation,” and typically refers to the presence of activated, pro-inflammatory microglia and astrocytes as well as infiltrating peripheral immune cells ^9,10^.

In humans, the presence of inflammatory microglia is associated with both aging and neurodegenerative conditions ^11,12^ and in mammals and *Drosophila*, increased innate immune signaling has been shown to induce or worsen neurodegeneration ^1,7,8,13^. *Drosophila* do not have microglia, but phagocytosis of neuronal processes and dead cells is carried out by other types of glia. The presence of uncleared cell corpses and debris is thought to lead to increased immune signaling ^14^ and glial phagocytic ability is known to decrease with age in both mammals and *Drosophila* ^15,16^. Thus, in aging, a lack of phagocytosis may contribute to increased neuroinflammation through the persistence of uncleared material that may in turn worsen neurodegeneration.

The humoral immune response in *Drosophila* is mediated primarily by two NFκB signaling pathways, Toll and Immune deficiency (Imd)^5,6,17^. These pathways activate transcription of antimicrobial peptides (AMPs) primarily in the fat body, a tissue with similarities to the vertebrate liver. However, there are additional complexities to these immune pathways. While AMPs are known to target bacteria and fungi, they have also been shown to contribute to neurodegeneration and other processes ^5,7^. Moreover, the Toll and Imd pathways activate transcription of other genes, are expressed in multiple tissues and act in concert with other pathways and phagocytic cells to remove invading pathogens ^5,17^.

The basic steps of phagocytosis are conserved across species ^18–20^. In *Drosophila*, Draper is a major phagocytic receptor with roles in phagocytosis of bacteria, debris and apoptotic cells throughout the body. In the brain, Draper has been shown to function in glia during the pruning of axons, clearance after axotomy, and in the removal of apoptotic cells ^21^. The absence of Draper in the brain leads to the persistence of neuronal corpses ^22–24^ and age-dependent neurodegeneration ^23,25^. However, the mechanisms by which the loss of Draper leads to neurodegeneration have yet to be determined.

Here we address whether inhibition of the cellular innate immune response, via mutants lacking the phagocytic receptor Draper, results in an overactivation of the humoral response. Specifically, we hypothesized that the absence of phagocytosis and presence of uncleared corpses would lead to increased innate immune signaling which would promote neurodegeneration. Indeed, we found elevated expression of anti-microbial peptides in the *draper* mutant, and found that suppression of the Imd pathway in either glia or fat body could reduce neurodegeneration in *draper* mutants. Taken together, these findings indicate that phagocytic defects lead to neurodegeneration through increased immune signaling, both systemically and locally in the brain.

## Results

### Aged *draper* mutants display dysregulation of anti-microbial peptide expression

To determine if innate immune signaling was affected in mutants lacking the phagocytic receptor Draper (*drpr*^*∆5*^), we extracted RNA from whole heads and performed RT-qPCR for three representative anti-microbial peptides: Attacin-A (*AttA)*, Attacin-D (*AttD)*, and Diptericin (*Dipt)*. We found that at an early age (5 days post-eclosion), *AttA* mRNA expression trended towards elevation in *drpr*^*∆5*^ heads, as compared to control heads (Figure 1A). At 14 days post-eclosion, this increase was more pronounced, with an average increase of around 700-fold as compared to 5-day old control flies. At 40 days post-eclosion, *AttA* expression in *drpr*^*∆5*^ mutants showed an increase of over 1500-fold as compared to 5-day old control flies. In contrast, 40-day old control flies showed an increase of about 14-fold compared to 5-day old flies. While *AttD* expression showed an increasing trend as *drpr*^*∆5*^ flies aged, this change was not significant (Figure 1B). *Dipt* expression similarly showed an increasing trend, but no significant change with age, and the increase appeared to be similar in control flies (Figure 1C). Taken together, these experiments suggest that innate immune signaling, particularly *AttA* expression, is drastically altered in *drpr*^*∆5*^ mutant heads. However, as this experiment involved extracting RNA from whole heads and not a specific tissue, we were not able to ascertain whether the observed dysregulated AMP expression could be attributed to the brain or another tissue in the head.

**Figure 1.**
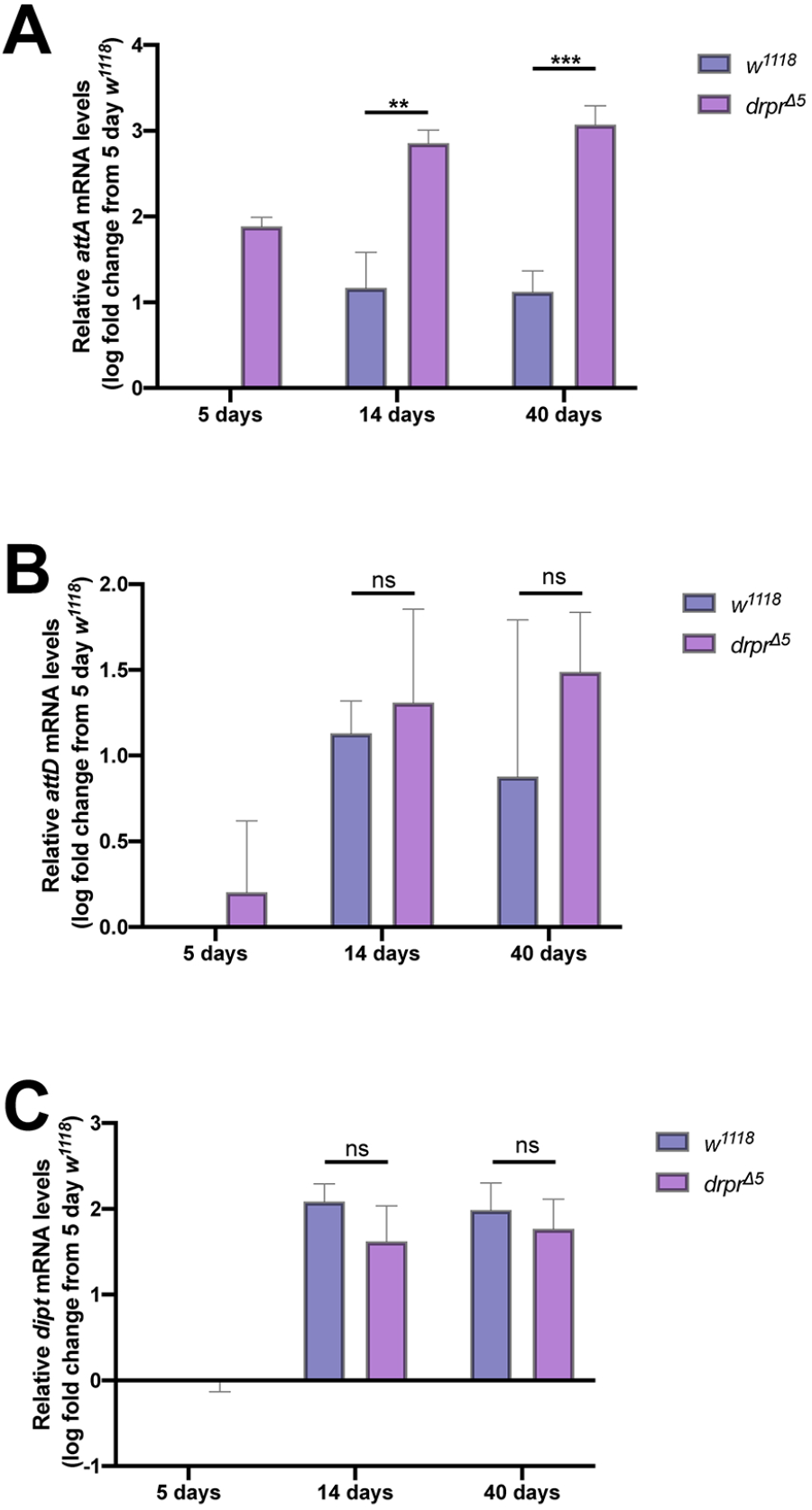
RT-qPCR shows dysregulation of innate immunity in *drpr*^*∆5*^ mutants. **A-C**. Fold change in expression of *AttA* (A), *AttD* (B), and *Dipt* (C) in whole heads of *drpr*^*∆5*^ or *w*^*1118*^ flies, as compared to 5-day old *w*^*1118*^, is shown. (\**p<0*.*05, **p<0*.*01, ***p<0*.*001, Unpaired t-tests with Welch correction were performed using Prism software)*.

### Attacin-GFP is expressed in neurons in young and aged, control and *draper* mutant brains at similar levels

To determine whether the dysregulation of AMP expression we observed in fly heads resulted specifically from brain expression, we used flies carrying the genetic reporter *AttGFP*, which contains the 2.4 kb regulatory region of *AttA* upstream of GFP and has been shown to report *AttA* expression ^26^. We first dissected brains of *AttGFP* flies and identified several cells in the brain that expressed *AttA* (Supplemental Figure 1A’-B’, arrowheads). Co-staining with anti-Repo antibody to mark glia (Supplemental Figure 1A”-B”’) revealed that these cells were not glia. Co-labeling with anti-Elav antibody to mark neurons (Supplemental Figure 1C-D”’) confirmed that most *AttGFP*-expressing cells were neurons. We then dissected young and aged *AttGFP* and *AttGFP, drpr*^*∆5*^ flies to determine whether the upregulation of *AttA* observed in *drpr*^*∆5*^ mutants through RT-qPCR resulted from brain expression. In young flies, *AttGFP, drpr*^*∆5*^ brains did not show a noticeable upregulation in *AttA* expression as compared to *AttGFP* brains (Figure 2A-B’). We then dissected the brains of flies aged to 40 days, and found once again that *AttA* expression was not noticeably different between *AttGFP* brains and *AttGFP, drpr*^*∆5*^ brains (Figure 2C-D’). Thus, we concluded that the increase in *AttA* expression in aged *drpr*^*∆5*^ mutants observed through RT-qPCR did not result from expression in the brain.

**Figure 2.**
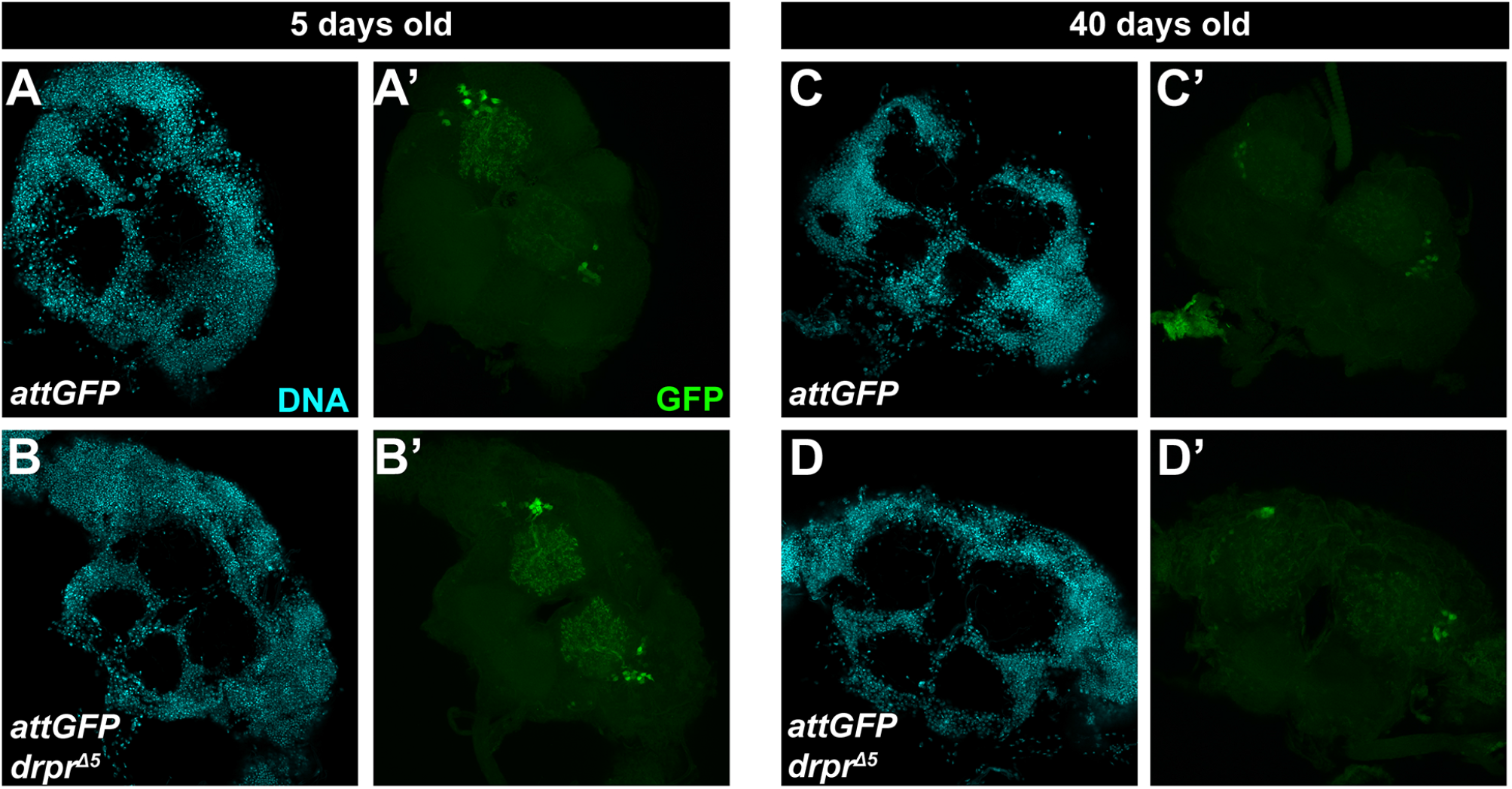
Brain expression of *AttA* is unchanged in aged *drpr*^*∆5*^mutants. **A-A’**. Representative images of 5-day old *AttGFP* brain stained with DAPI (A) and anti-GFP (A’). **B-B’**. Representative images of 5-day old *AttGFP, drpr*^*∆5*^ brain stained with DAPI (B) and anti-GFP (B’). **C-D’**. Representative images of 40-day old *AttGFP* (C-C’) or *AttGFP, drpr*^*∆5*^ (D-D’) brain stained with DAPI (C, D) and anti-GFP (C’,D’).

### Attacin-GFP expression is increased in the aging *drpr* mutant fat body

To visualize *AttA* expression in tissues other than the brain, we cryosectioned whole heads of *AttGFP* and *AttGFP, drpr*^*∆5*^ flies. We used tissue morphology, nuclear labeling by DAPI, and anatomical location to identify fat body tissue, brain tissue, and eyes in head sections (Figure 3B-B’). The fat body was a tissue of particular interest, as it is associated with immune signaling. We used BODIPY, which labels lipids (Figure 3A), to distinguish the pericerebral fat body from other tissues surrounding the brain. As observed in the dissection of whole brains (Figure 2), there was no difference in *AttGFP* expression in the brain between any groups observed in cryosections (Figure 3C-F). We thus turned our attention to the pericerebral fat body. We found that in young flies (1-2 days post-eclosion), there was no difference in *AttA* expression between *AttGFP* (control) flies (Figure 3C) and *AttGFP; drpr*^*∆5*^ flies (Figure 3D). However, in aged flies, both *AttGFP* (Figure 3E) and *AttGFP; drpr*^*∆5*^ (Figure 3F) showed an increase in *AttGFP* expression in the pericerebral fat body. While this increase was present in some of the sections from *AttGFP* flies, it was present in all sections from *AttGFP; drpr*^*∆5*^ (n=7 flies). The percentage of GFP-positive pericerebral fat body was quantified in several sections per head, and we found a highly significant increase in *AttA* expression in aged, but not young, *drpr* mutants, compared to age-matched controls (Figure 3G).

**Figure 3.**
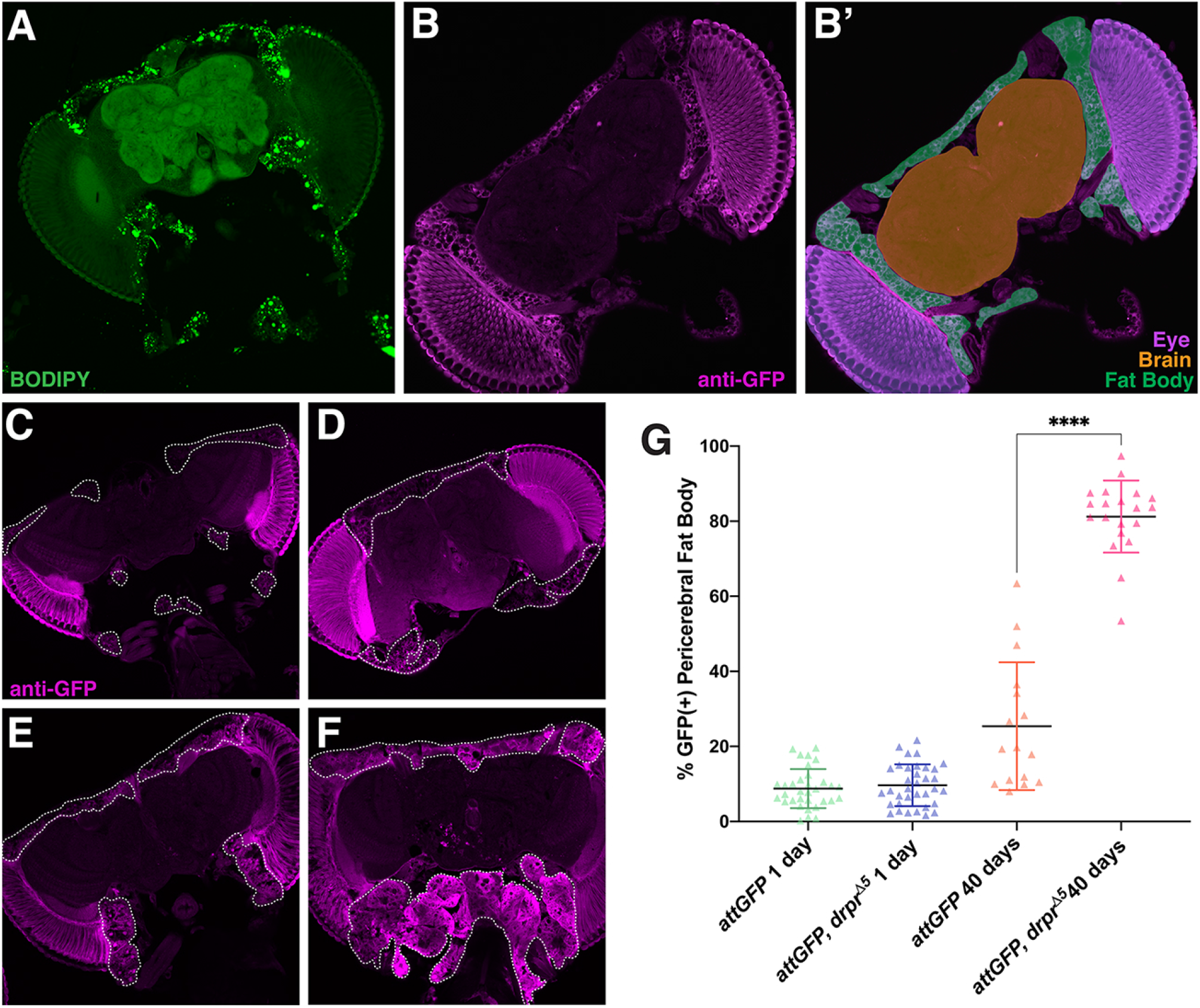
Pericerebral fat body expression of *AttA* is increased in aged *drpr* mutants. **A**. BODIPY staining (lipids) was used for positive identification of fat body cells in cryosectioned *Drosophila* heads. **B-B’**. Cryosection of *AttGFP* fly head stained with anti-GFP (magenta, B), with brain, eyes, and fat body highlighted in orange, purple, and green, respectively (B’). **C-D**. Representative sections of young *AttGFP* (C) and *AttGFP; drpr*^*∆5*^ (D) heads, stained with anti-GFP (magenta), showing no difference in *AttA* expression. Dotted white line surrounds regions of fat body. **E-F**. Representative sections of aged *AttGFP* (E) and *AttGFP; drpr*^*∆5*^ (F), stained with anti-GFP (magenta), showing increased *AttA* expression in *drpr*^*∆5*^pericerebral fat body. **G**. Quantification of GFP-positive pericerebral fat body in young and aged *AttGFP* and *AttGFP; drpr*^*∆5*^ flies. Values shown are mean ± S.D. \*\*\*\**p<0*.*0001 (unpaired t-tests were performed using Prism software)*. Sections were quantified from 3 replicate experiments (*AttGFP* young n= 8, *AttGFP; drpr*^*∆5*^ young n= 9, *AttGFP* aged = 10, *AttGFP; drpr*^*∆5*^ n=7).

We next determined whether the increase in *AttA* expression was specific to the pericerebral fat body or if it was a systemic effect. We cryosectioned whole bodies of *AttGFP* and *AttGFP, drpr*^*∆5*^ flies and stained with anti-GFP antibody to visualize *AttA* expression. We used cell morphology and anatomical location to identify fat body cells in the abdomen. As with the pericerebral fat body, expression of *AttA* in the abdominal fat body was unchanged between the young *AttGFP* control flies (Supplemental Figure 2A-A’’’) and *AttGFP, drpr*^*∆5*^ flies (Supplemental Figure 2B-B’’). In 40-day old flies, there was an increase in *AttA* expression between *AttGFP* control flies (Supplemental Figure 2C-C’’) and *AttGFP, drpr*^*∆5*^ flies (Supplemental Figure 2D-D’’). However, this increase was not consistent across all samples where some abdomens did not show an increase in *AttA* expression (Supplemental Figure 2E-E’’). This suggests that the upregulation of immune signaling was more pronounced in the head of *drpr* mutants.

### Glial and fat-body, but not neuronal, knockdown of *Relish* in *draper* mutants attenuates neurodegeneration

To determine whether the neurodegeneration observed in *drpr*^*∆5*^ mutants is mediated by an overactive immune response, we genetically inhibited the Imd pathway in neurons, glia, or fat body cells in a *drpr*^*∆5*^ mutant background by expressing dsRNA targeting the NFκB transcription factor *Relish* using Gal4 drivers specific to these cell types. We then aged the flies and performed cryosectioning to measure levels of vacuolization, a hallmark of neurodegeneration in *Drosophila*. We found, as expected, that *drpr*^*∆5*^ mutants carrying only the GAL4 drivers showed high levels of neurodegeneration compared to controls (Figure 4A-F, J). *drpr*^*∆5*^ mutants expressing *Relish* RNAi in neurons with *elav-Gal4* did not show decreased levels of neurodegeneration (Figure 4I, J). *drpr*^*∆5*^ mutants carrying only the fat body driver *ppl-GAL4* showed high levels of neurodegeneration but *drpr*^*∆5*^ mutants expressing *Relish* RNAi in fat body showed significantly less vacuolization (Figure 4E, H, J). Similarly, inhibiting *Relish* in glia with *repo-GAL4* in a *drpr*^*∆5*^ mutant background significantly reduced the level of vacuolization, as compared to *drpr*^*∆5*^ mutants carrying only the glial driver (Figure 4 D, G, J). Taken together, these findings indicate that activation of the Imd pathway in glia and fat body, but not neurons, contributes to neurodegeneration seen in the *drpr*^*∆5*^ mutant.

**Figure 4.**
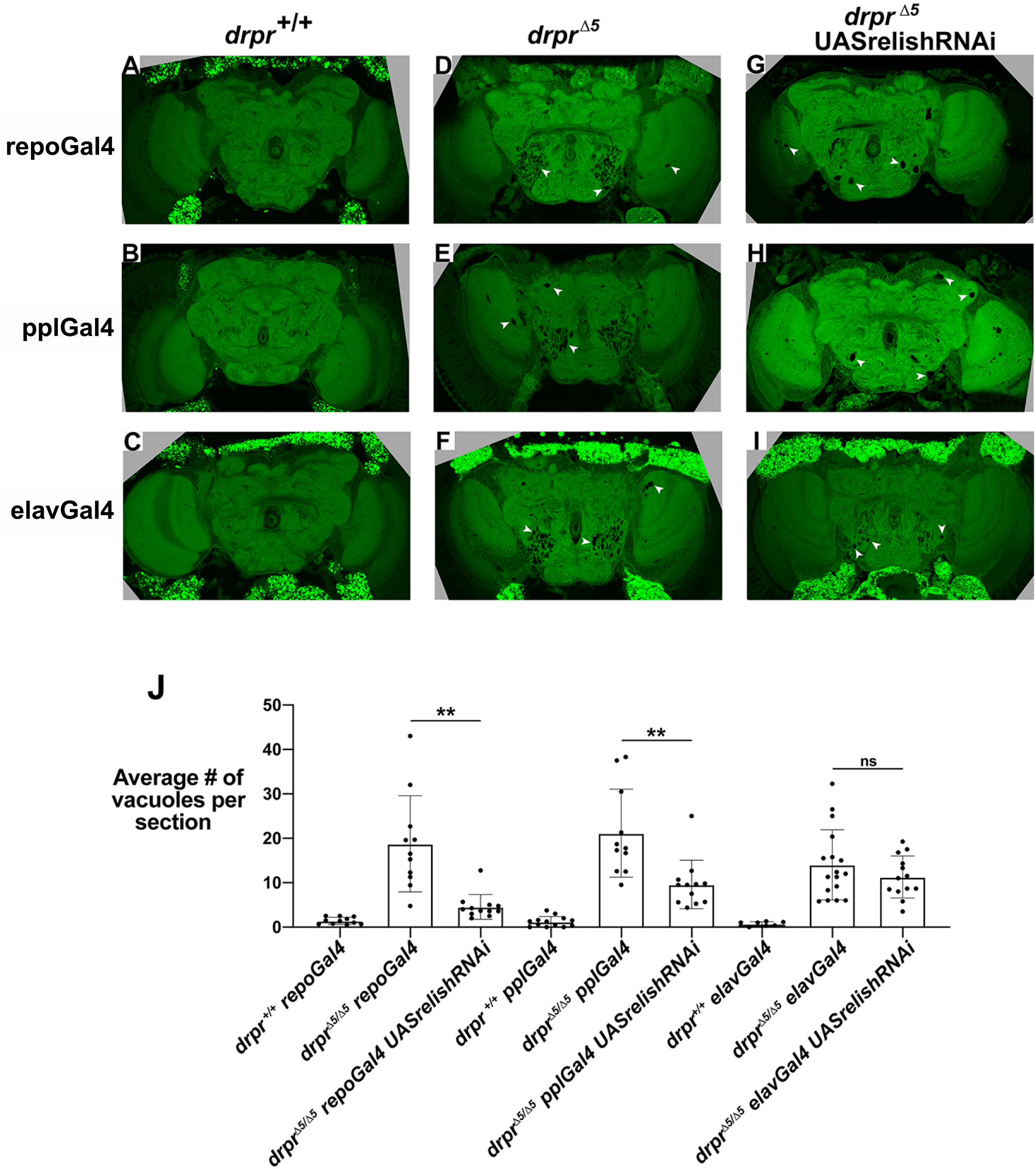
Knockdown of *Relish* in glia and fat body, but not neurons, attenuates neurodegeneration in *drpr* mutant flies. **A-C**. Representative sections of 40-day old *drpr*^*+/+*^ flies carrying the glial driver *repo-Gal4* (A), the fat body driver *ppl-Gal4* (B), or the neuronal driver *elav-Gal4*, stained with BODIPY. **D-F**. Representative sections of 40-day old *drpr*^*∆5*^ flies carrying *repo-Gal4* (D), *ppl-Gal4* (E), or *elav-Gal4* (F) stained with BODIPY. White arrowheads point to vacuolar lesions. **G-I**. Representative sections of 40-day old *drpr*^*∆5*^ flies with *Relish-RNAi* expressed in glia (G), fat body (H), or neurons (I). White arrowheads point to vacuolar lesions. **J**. Average number of vacuolar lesions per section for each genotype. Each dot represents one head, with at least two sections analyzed per head. (\*\**p<0*.*01. Brown-Forsythe and Welch ANOVA and unpaired t tests with Welch correction were performed using Prism software. P-values shown for unpaired t tests with Welch correction)*

## Discussion

Chronic activation of innate immunity in the brain is known to play a role in neurodegenerative disease progression, although the mechanisms underlying this role remain unclear ^27^. Microglia, the resident macrophages of the mammalian brain, can become “activated”, releasing pro-inflammatory cytokines and reactive oxygen species, damaging nearby neurons ^28^. However, “activated” microglia are also thought to be protective at times, clearing dead cells, cellular debris, and disease-associated protein aggregates ^29,30^. Interestingly, in aging, the phagocytic abilities of microglia decrease, while their pro-inflammatory reactivity increases ^16^. This interplay between phagocytosis and innate immunity in the brain remains poorly understood. In *Drosophila*, recent findings have suggested that increased innate immune signaling in the brain is similarly detrimental ^7,8,13^. As we and others have shown, phagocytic defects in *Drosophila* glia lead to age-dependent neurodegeneration ^23,25^. Thus, *Drosophila* provide a useful model in which to untangle the links between glial phagocytosis, innate immune signaling, and neurodegeneration.

As increased innate immune signaling is thought to cause or worsen neurodegeneration, and the presence of uncleared cell corpses and debris is thought to lead to activation of innate immunity, we hypothesized that the neurodegeneration observed in *drpr*^*∆5*^ flies could be caused by overactive innate immune signaling. Thus, we first measured AMP expression as a readout for the Imd pathway and found that *AttA* was upregulated in the aged *drpr*^*∆5*^ mutant head. We then used cryosectioning to identify which tissue was responsible for the elevated *AttA* expression and found that that this upregulation was primarily the result of expression in the fat body. To determine whether this overactive immunity was contributing to neurodegeneration, we knocked down *Relish* in the fat body, glia, and neurons in the *drpr*^*∆5*^ mutant and measured neurodegeneration. We found that inhibiting the Imd pathway in glia and fat body, but not neurons, partially suppressed neurodegeneration in the *drpr*^*∆5*^ mutant. These findings suggest that increased innate immune activity, both systemically and in glia, drive neurodegeneration in the *drpr*^*∆5*^ mutant.

A previous study found that increased activation of the Imd pathway can induce age-dependent neurodegeneration ^7^. Mutations in the *dnr1* (*defense repressor 1*) gene, a negative regulator of the Imd pathway, result in increased immune signaling and neurodegeneration. The authors also induced AMP expression in the brain, both genetically and by bacterial injection, and found this sufficient to cause neurodegeneration ^7^. Interestingly, when the authors activated systemic innate immunity by injecting bacteria into the thorax, they did not see neurodegeneration. The authors state that this is because AMPs are not expected to cross the blood-brain barrier. Our finding that suppressing systemic immunity is sufficient to attenuate neurodegeneration suggests the possibility that some AMPs do cross the blood-brain barrier, or that systemic innate immunity can be neurotoxic in other ways. It is possible that the *drpr*^*∆5*^ mutant has increased blood-brain barrier permeability, although to our knowledge this has not been investigated. Due to the nature of the *AttGFP* reporter, we were only able to visualize transcriptional regulation of *AttA*, rather than localization of the peptide.

In humans, systemic inflammation can occur chronically or acutely, driven by underlying causes such as infection, chronic illness, and aging. Cytokines and other pro-inflammatory mediators can affect the brain by crossing the blood-brain barrier or entering the circumventricular organs ^31,32^. Systemic inflammation is thought to induce activation of glia, leading to inflammation in the CNS ^33^. In aged or diseased brains, glia are thought to be primed in a way that makes them more reactive to stimuli capable of inducing a pro-inflammatory state ^34^. Chronic, low-level systemic inflammation is also thought to play an important role in neurodegenerative disease. Evidence for this comes from long-term studies showing that factors known to increase levels of inflammation, such as smoking, obesity, vascular disease, and periodontal disease increase the risk of neurodegenerative disease ^35,36^. Additionally, long term use of NSAIDs reduces the risk of AD, suggesting that maintaining low levels of systemic inflammation earlier in life can be protective ^37^.

Our study identifies a source of systemic inflammation in *Drosophila*: defective phagocytosis. Whether this results from the persistence of apoptotic corpses, debris, pathogens, or another mechanism remains unclear. Our finding that inhibition of immune signaling in glia attenuated neurodegeneration suggests that elevated systemic immune signaling may drive neurodegeneration through activation of glia, as is thought to be the case in mammals. In *Drosophila*, activation of glia has primarily been described in the context of phagocytic clearance of injured axons ^38^. Whether glia engage in other pro-inflammatory signaling remains unclear, although Imd signaling has been reported in glia ^7,39^. While we did not see elevated expression of *AttA* in glia in the *drpr*^*∆5*^ mutant, *Relish* regulates the expression of many genes including other AMPs ^17^. Identification of these key transcriptional targets of Relish will provide insight into how local and systemic inflammation drives neuroinflammation and neurodegeneration in *Drosophila*.

### Materials and Methods

### Fly Stocks and husbandry

The following fly strains were used: *drpr*^*∆5*^ *FRT2A* ^40,41^, *UAS-Relish RNAi* (Vienna *Drosophila* Stock Center #49414), *AttGFP* (^26^ provided by Dr. Mitch Dushay), *repo-GAL4/TM3* (Bloomington *Drosophila* Stock Center (BDSC) #7415), *elav-GAL4* (BDSC #458), *ppl-GAL4* (^42^ RRID:BDSC_58768, provided by Dr. Michael O’Connor) and *w*^*1118*^ (BDSC). All fly lines were raised at 25°C on standard cornmeal and molasses food.

### Dissection and Histology

Flies were anesthetized on CO_2_ pads before dissection. Brains were dissected out of the cuticle in 4% paraformaldehyde (PFA) in phosphate-buffered saline with 0.1% Triton-X 100 (PBT) and transferred to 0.5 ml centrifuge tube with fresh dissection media. Whole heads were removed from bodies in fixation media (4% PFA in PBT). The proboscis was then removed, with care being taken not to damage the head. Heads were then placed in fresh fixation media and allowed to rotate for 2-4 hours at room temperature. For whole body preparations, the wings, legs, and proboscis were removed and the abdomen was lightly punctured in fixation media (4% PFA in PBT). Flies were then placed in fresh fixation media and incubated at room temperature with rotation for 4 hours.

For cryosectioning after dissection, PFA was removed and replaced with PBT for up to an hour. PBT was then removed and replaced with 15% sucrose in PBT. Heads or bodies were incubated for 48 hours in sucrose at 4°C with rotation. 15% sucrose was then replaced with 30% sucrose in PBT, and tissues were incubated for a further 48 hours at 4°C with rotation. Tissues were then embedded in optimal cutting temperature (OCT) media (Fisher #23-730-571) in aluminum foil molds and frozen on powdered dry ice, before being stored at -80°C until sectioning. Tissues were sectioned at -19 to -21°C, with heads at a thickness of 18 µm and whole bodies at 25µm unless noted otherwise.

### Staining procedures

For whole mount staining, dissected brains were incubated for 50 minutes in fresh fixation media at room temperature. Samples were then permeabilized in three washes for a total of up to one hour in 1X PBT at room temperature. Samples for antibody staining were incubated in PBANG (PBT, bovine serum albumin (BSA), and normal goat serum (NGS)) for one hour at room temperature or overnight at 4°C. Samples were then incubated with primary antibody diluted in PBANG for 4-5 days at 4°C with rotation. Samples were then washed in PBT for a total of one hour, before being incubated in secondary antibody diluted in PBANG for 4 hours at room temperature or two days at 4°C in the dark with rotation. Samples were then transferred to Vectashield with DAPI (Vector Laboratories) and stored in the dark at 4°C until mounting on slides.

Sections on slides were washed in coplin staining jars in PBS before being incubated for two hours in primary antibody diluted in PBS. Slides were then washed twice in PBS for up to one hour before being incubated for 45 minutes in secondary antibody diluted in PBS. Slides were then again washed at least three times in PBS for a total of one hour. Coverslips were then mounted with two drops of Vectashield with DAPI (Vector Laboratories), then sealed with nail polish.

The following primary antibodies were used: Anti-GFP (Torrey Pines Biolabs Cat# TP401 071519, RRID:AB_10013661, 1:100), Anti-Elav (DSHB Cat# Elav-9F8A9, RRID:AB_528217,1:100) and Anti-Repo (DSHB Cat# 8D12 anti-Repo, RRID:AB_528448, 1:100). Secondary antibodies were obtained from Jackson Laboratories and were used at 1:200 or 1:400.

For BODIPY (lipid) staining, whole tissues were dissected, fixed, and permeabilized as described above, and were incubated at room temperature in the dark in a solution of 1:1000 of 1mg/ml BODIPY (Invitrogen #D3922) diluted in DMSO:1X PBS for 2 hours. Samples were then washed three times in PBS and stored in Vectashield with DAPI (Vector Laboratories) until coverslipping. After sectioning and washing in PBS as described above, sections on slides were incubated in a solution of 1:1000 of 1mg/ml BODIPY (Invitrogen #D3922) diluted in DMSO:1X PBS for 45 minutes at room temperature or overnight at 4°C, protected from light. Slides were then washed three times with 1X PBS, then a coverslip was added as described above.

### Microscopy

Samples were visualized by either fluorescence and light microscopy (Olympus BX-60), or confocal microscopy (Olympus FluoView FV10i, or Nikon C2Si). Briefly, image analysis was performed using ImageJ. Image processing for figure production was performed using Adobe Photoshop or Adobe Illustrator.

### Image processing and quantification

To measure Attacin-GFP expression in the fat body, the fat body area was first defined using cell morphology, anatomy, and DAPI. The region of interest was outlined in FIJI. A minimum threshold was determined using the center areas of the brain as a measure for background. The % of the fat body area which was above the threshold was then measured and recorded. Sections which were folded, improperly stained, or overly damaged were discarded.

To measure neurodegeneration, heads were cryosectioned and stained with BODIPY as described above. Sections were imaged by confocal microscopy as outlined above. Images were converted to PNG files in FIJI, then opened in Adobe Photoshop for vacuole counting. Vacuoles larger than four microns in diameter were counted, using the brush tool as a size guide. At least two sections per brain were analyzed. Sections with most of the central brain and optic lobe were selected for counting, and sequential sections were not used, to avoid counting the same vacuoles twice. Researchers were blinded to genotype. Vacuoles located in the retina were not counted. Any cracks in the tissue were not counted, as these result from processing.

### RT-PCR

Flies were aged in groups of 15 males and 15 females until the appropriate age was reached. Flies were then placed in cryovials and snap-frozen in liquid nitrogen in groups of ∼30. Flies were stored at -80°C until ready for use, or processed immediately. Frozen flies were decapitated by vortexing for 10 seconds, placing back in liquid nitrogen, and then repeating the process one more time. Heads were separated from bodies rapidly using a paintbrush, over liquid nitrogen. Heads were then homogenized in QIAlysis solution. RNA was then extracted using the Qiagen RNeasy kit. The final concentration of RNA from this process was about 80 ng/µl for 30 heads. RNA was stored at -80°C until cDNA synthesis.

The Maxima First Strand cDNA Synthesis Kit for RT-qPCR (Thermo Scientific #K1671) was used to produce cDNA. Kit components and samples were thawed on ice, and RNA concentration was determined using a NanoDrop. Sterile, RNase-free tubes on ice were used for reactions. To each tube, the following were added: 1 µl of 10X dsDNase buffer, 1 µl of dsDNase, 0.1 µg of total RNA, and nuclease-free water to reach 10 µl. Reactions were gently mixed and briefly centrifuged on a table-top centrifuge. Reactions were then incubated at 37°C for 2 minutes, then chilled on ice. The following components were then added to each tube: 4 µl of 5X Reaction Mix, 2 µl of Maxima Enzyme Mix, and 4 µl of nuclease-free water. Reactions were gently mixed and briefly centrifuged on a table-top centrifuge. Reactions were then incubated for 10 minutes at 25°C, followed by 15 minutes at 50°C, then 5 minutes at 85°C. The reaction products were then immediately used for RT-qPCR.

The GoTaq qPCR Master Mix (Promega, A6001) was used to perform RT-qPCR. cDNA was diluted to 5 ng/µl, and primers diluted to 10 mM. Kit components were thawed on ice. In nuclease-free tubes on ice, master mixes were prepared for each primer. In a 384-well plate, 2 µl of cDNA was loaded into each well (except NTC, in which water was added instead). 8 µl of Master Mix for the appropriate primer set was then added to each well. The plate was sealed with a clear plate cover, then briefly centrifuged. Samples were then analyzed in an ABI 7900ht real time PCR machine.

### Statistical analysis

All statistical analyses were performed using Prism software. For RT-qPCR, unpaired t-tests with Welch correction were performed on log-transformed delta-delta-CT values and samples were normalized with Rpl32. For determining the % of fat body positive for GFP, unpaired t-tests were performed. For measuring neurodegeneration using cryosectioning, Brown-Forsythe and Welch ANOVA and unpaired t tests with Welch correction were performed.

## Acknowledgements

We thank Mel Feany, Susan Holmes, Angela Ho, Tuan Leng Tay and members of our laboratory for helpful discussions and comments on the manuscript. We thank JiSoo Park for assistance with experiments. We thank the Vienna and Bloomington *Drosophila* Stock Centers, Mitch Dushay, and Michael O’Connor for generously providing fly strains, and the Developmental Studies Hybridoma Bank for antibodies. We thank our funding sources: NIH Grants R21 AG056158 and R35 GM127338 to KM, the Beckman Foundation to KT and UROP at Boston University to LD.

## Author contributions

Conceptualization: JEE and KM; Methodology: JEE; Investigation: JEE, GL, HG, KT and LD; Writing: JEE and KM; Funding Acquisition: KM, Supervision: KM.

---

**Supplemental Figure 1.**
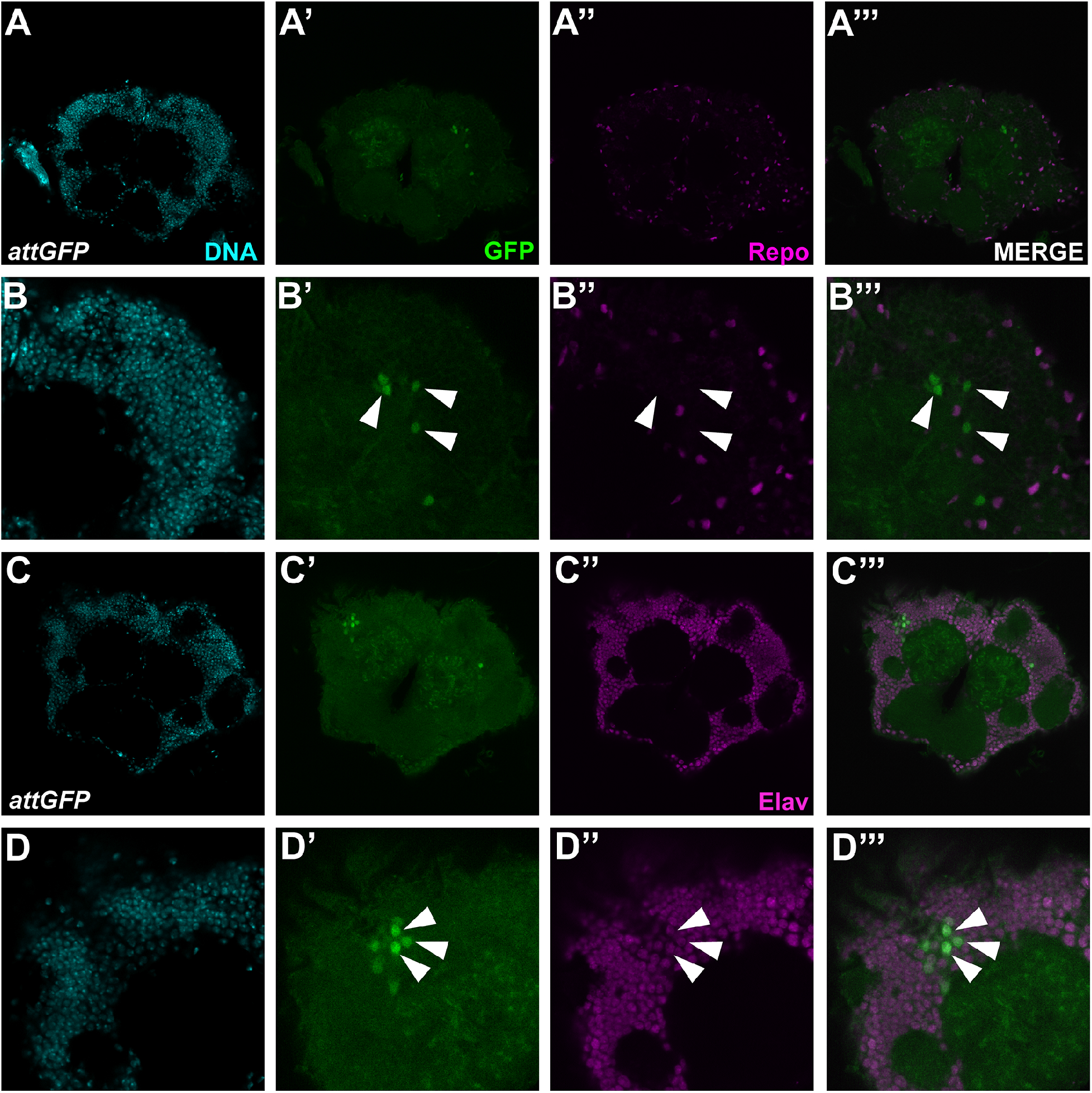
*AttA* is expressed in select neurons. A-A’’’. Representative images of *AttGFP* brain stained with DAPI (A), anti-GFP (A’), and anti-Repo (A’’). A’’’ shows a merged image of anti-GFP and anti-Repo. **B-B’’’**. Enlargements of images in A-A’’’. White arrowheads point to GFP-positive cells. **C-C’’’**. Representative images of *AttGFP* brain stained with DAPI (C), anti-GFP (C’), and anti-Elav (C’’). C’’’ shows a merged image of anti-GFP and anti-Elav. **D-D’’’**. Enlargements of images C-C’’’. White arrowheads point to GFP-positive cells.

**Supplemental Figure 2.**
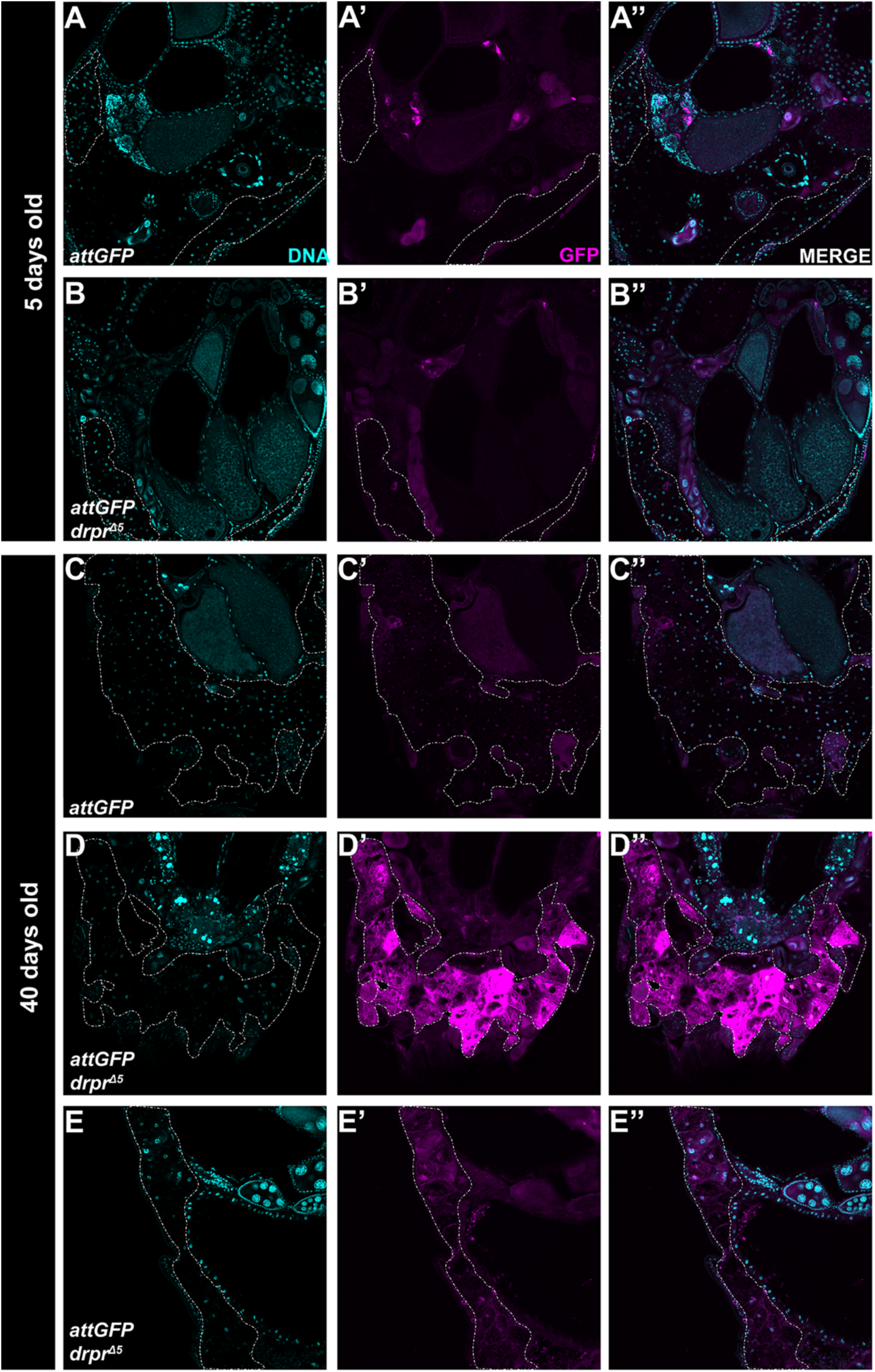
Abdominal fat body expression of *AttA* is increased in some aged *drpr* mutants. A-B’’. Representative images of cryosectioned abdomens from 5-day old *AttGFP* (A-A’’) and *AttGFP, drpr*^*∆5*^ (B-B’’) flies stained with DAPI and anti-GFP. Dotted line outlines fat body. **C-E’’**. Representative images of cryosectioned abdomens from 40-day old *AttGFP* (C-C’’) and *AttGFP, drpr*^*∆5*^ (D-E’’) flies stained with DAPI and anti-GFP. Dotted line outlines fat body. Abdomen in D shows strong expression but E shows more moderate levels.

